# Make it so: Rapid and affordable plasmid sequencing on ONT platforms with PICARD-seq

**DOI:** 10.64898/2026.09.19.752805

**Authors:** Rebecca Wolters, Johannes W. Debler, Gabriella Go, Anna Bond, Florence Ly, Cody Ryall, Jed Orrock, Nicolas Tabanera Valle, Bryony Hodge, Farley M Kwok van der Giezen, Patrick Gong, Harish Jadhav, Mojtaba Bagherian, Zakia Thevarajoo, Adil Khan, Gabrielle Herring, Jia Yuan Zhu, Tessa Swain, John Blinco, Faiza Chowdhury, Christian Pflueger, Suruchi Roychoudhry, Asher Floyd, Ryan Lister, Archa Fox, James P B Lloyd

## Abstract

Plasmid construction underpins molecular biology and synthetic biology, yet validation is often limited to the inserted fragment rather than the whole plasmid, and around a third of laboratory-made plasmids carry sequence errors that can affect function. Sanger sequencing scales poorly across whole plasmids, while short-read approaches cannot resolve the repeated DNA parts, such as promoters, that are common in synthetic constructs. We present PICARD-seq, a rapid nanopore-based protocol that uses off-the-shelf Tn5 rapid barcoding reagents and a MinION to sequence pools of whole plasmids in under a day, and we systematically benchmark the computational pipelines available for analysing the resulting data. Using a curated set of 25 plasmids of known sequence spanning 3.0-20.6 kbp, including various dilution series and repetitive multi-part constructs, we ran five independent replicates of each pipeline. The ONT EPI2ME Clone Validation workflow was fast (13-18 min) but stochastic, varying between replicates for both plasmids assembled and what sequence was returned; Canu outperformed the default Flye assembler, and reducing the minimum coverage parameter from 60x to 20x substantially improved assembly of large, repetitive, and dilute samples. The ensemble assembler Autocycler was slower (81-111 min) but gave the highest and most consistent rate of recovering the expected sequence. Complementary read mapping with minimap2 distinguished genuine sequence differences from assembly artefacts. Applying PICARD-seq to problematic plasmids revealed backbone concatemers, a misincorporated promoter part, and a mixed population of rearranged molecules in a repetitive construct. PICARD-seq makes routine whole-plasmid validation practical and affordable for individual laboratories.

## Introduction

Synthetic biology holds much promise for advancing various aspects of biotechnology from drug production, bioremediation, and agriculture (Rylott and Bruce, 2020; Yan et al., 2023; Roychoudhry et al., 2026). However, much of this is built upon standard molecular cloning protocols and validating the successful construction of plasmids. It is common in many molecular biology laboratories to only sequence the portion of DNA inserted into a plasmid rather than the whole plasmid, including the backbone. Furthermore, it is widespread practice to only sequence after a PCR or DNA synthesis step, as these are the most error prone steps in the process. However, there are many cases where mutations or assembly errors may occur independent of PCR or synthesis. These errors can also occur in the backbone of the plasmid rather than in the insert and it is known that the properties of the backbone can influence the performance of the plasmid in bacteria (Tas et al., 2021). For instance, Bai and colleagues found that 35% of examined lab-made plasmids contained sequence errors that could affect their function to varying degrees (Bai et al., 2025), highlighting that this is a prevalent problem.

Ongoing advances in DNA synthesis technologies and synthetic biology methods are pushing the length of plasmids to new large sizes and levels of complexity and repetitiveness (Robinson et al., 2026), often with repeated units because of limited characterised parts for particular organisms. Consequently, it is very important to be able to ensure the plasmid is as expected before testing in a biological context. Sanger sequencing has been the gold standard for sequencing plasmids for decades. However, this requires the use of a primer to define the start of the sequencing read. This lends itself well to the sequencing of inserts after a specific cloning step, but not towards the sequencing of the whole plasmid, including the backbone. Given the average length from Sanger sequencing is only around 900 bp, the sequencing of a large plasmid would require many sequencing reads which can be cost prohibitive. For example, to fully sequence a 15 kbp plasmid, at least 17 sequencing reactions would be expected, with well placed primer start sites for each read to sequence the whole plasmid. Assuming $10 Australian dollars (AUD) per sequencing reaction ($7.12 United States dollars; USD), this would bring the total cost to $170 AUD ($121.09 USD), which is prohibitively expensive, and does not even include the primer design and synthesis costs.

Multiple protocols for cost-effective high throughput sequencing of whole plasmids have been published for Illumina sequencing platforms (Shapland et al., 2015; Gao et al., 2021; Suzuki et al., 2026). However, they are limited by high capital investment for the equipment needed, such as a MiSeq, and they are innately limited by the short read length. If a repeated DNA part of the plasmid is larger than the read length used for sequencing then it will be very challenging to assemble the plasmid as a single contig *in silico*.

Oxford Nanopore Technologies (ONT) workflows have been used to sequence whole plasmids for clinical purposes, often as a single plasmid using an entire flow cell, which is prohibitively expensive for routine sequencing (Brown et al., 2023). ONT has also been used to sequence plasmids as multiplexed reactions to save on costs and increase throughput (Emiliani et al., 2022; Mumm et al., 2023; De Oliveira et al., 2026; Schimke and Vollmers, 2026). Here, we present our approach for rapid sequencing of synthetic plasmids using an ONT platform called PICARD-seq (Lloyd et al., 2026) and the computational analysis of the data generated. We have compared various computational pipelines to determine the best way to analyse the data. Our protocol uses standard off the shelf reagents from ONT from their rapid library prep, standard flow cell priming and wash kits, combined into one simple to follow protocol in the lab (Lloyd et al., 2026). The affordability of a MinION, suitable computer to run it, and easy to acquire reagents means that this protocol is highly accessible by many researchers, relative to other approaches, such as those that require larger sequencing machines with higher costs associated with initial purchases.

## Results

### Overview of the PICARD-seq workflow

Using the PICARD-seq protocol (Lloyd et al., 2026), a pool of plasmids can be sequenced in 0.5 to 2 hours on a MinION (Figure 1), which is preceded by less than one hour of hands-on time to prepare the pooled library of plasmids for sequencing, and to prime the flow cell. To enable reuse of the same flow cell, additional time is needed to perform a wash of the flow cell with a nuclease solution to prepare it for future runs. A fresh flow cell can thereby be used and re-used by repeating the wash and priming cycles between runs. Alternatively, old flow cells that have been partially used for other purposes such as whole gene sequencing can be used, and is an effective cost saving measure. We have recently reported a detailed step-by-step protocol for the experimental components of the PICARD-seq workflow (Lloyd et al., 2026). To assess the utility of PICARD-seq across a range of plasmids, and to establish a reliable computational pipeline for the analysis of the data, we sequenced a diverse set of plasmids used in synthetic biology, as described in Table 1. Computational analysis of the plasmids is usually quick, taking less than an hour with most pipelines. Therefore, the whole workflow can be run in less than a day, allowing for fast return of results to eager researchers, and significant acceleration of research progress.

**Figure 1:**
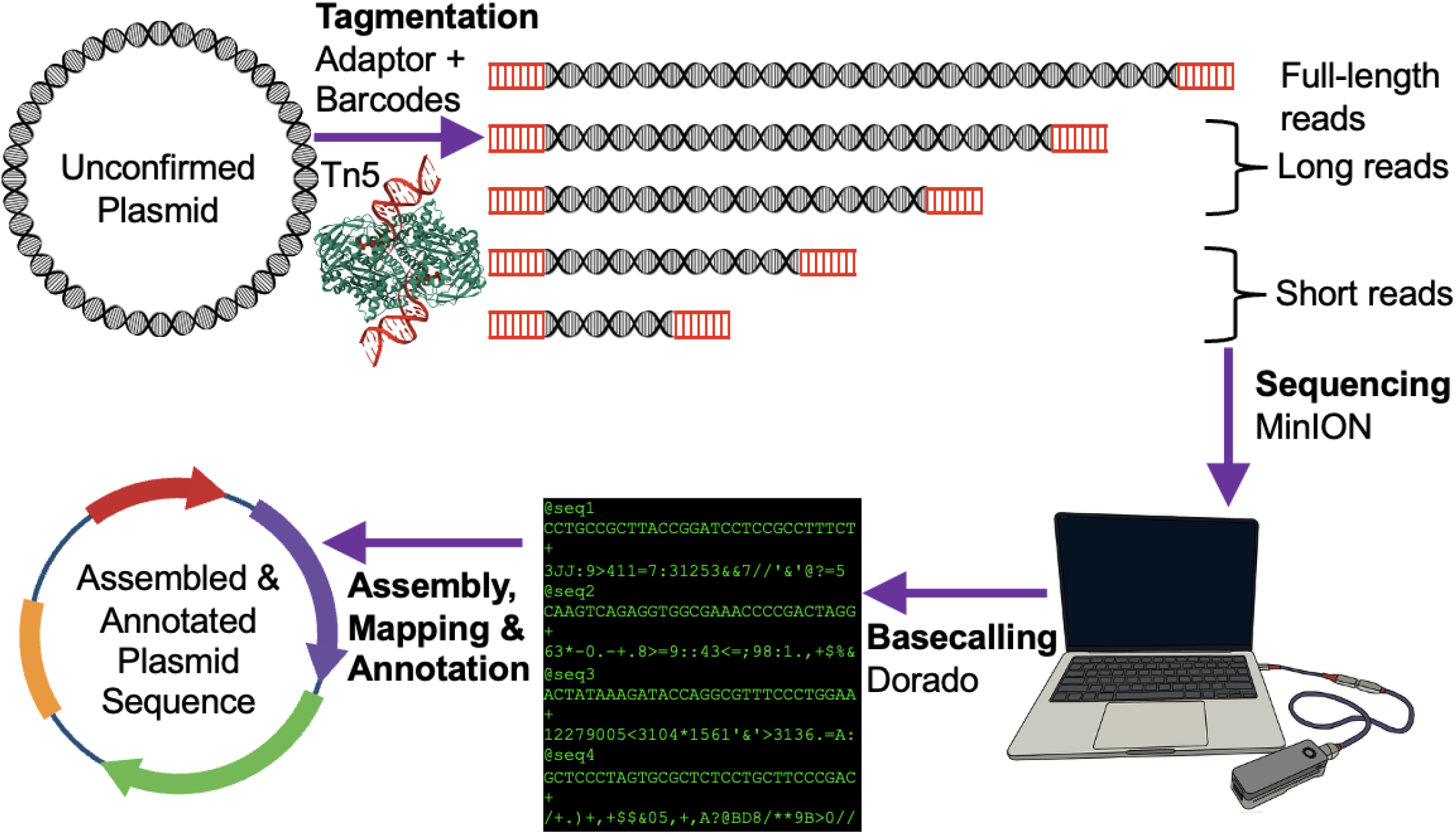
Overview of PICARD-seq pipeline. Plasmid DNA is tagmented with Tn5 preloaded with adaptor and barcode sequences from ONT. DNA fragments of different lengths are typically generated, reflecting fragments of full plasmid length (a single cut by Tn5 to linearize the plasmid), long reads (but not full-length, cut multiple times by Tn5) and short reads (<1 kb reads), all of which are sequenced on an ONT MinION. Dorado software (ONT) then converts the raw electrical signals (POD5 files) of library molecules into its corresponding predicted DNA sequence (FASTQ files). Sequenced reads can be assembled into a *de novo* assembled plasmid, automatically annotated, and compared to expected DNA sequences.

**Table 1:**
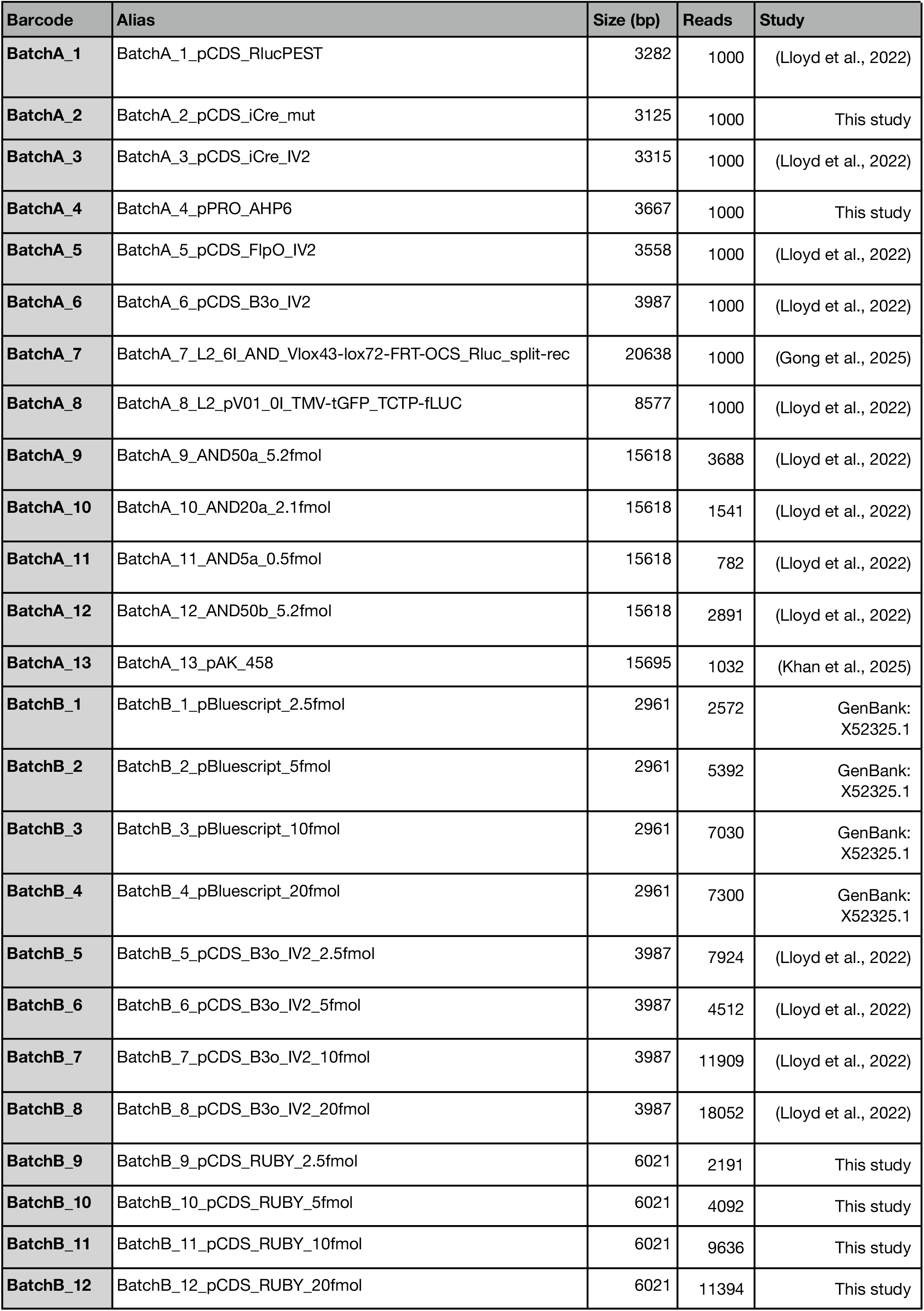
Test plasmids for benchmarking computational pipelines used to assemble plasmids from PICARD-seq.

### EPI2ME Clone Validation workflow is fast but displays stochasticity

ONT has provided a Nextflow computational workflow to analyse their data via their EPI2ME software. The EPI2ME Clone Validation workflow can take advantage of two different DNA assemblers within EPI2ME: Flye (default) and Canu (Koren et al., 2017; Kolmogorov et al., 2019). We set out to test the EPI2ME Clone Validation workflow with both of these assemblers. We have curated a set of 25 plasmid samples that have known sequences as established through various prior testing (Table 1). The expected sequence should have a 100% match to the assembled sequence. In some cases, there are repeated plasmids that have been diluted to different levels to test the effects of input DNA level on the sequencing reaction (Table 1). In preliminary testing, we noticed that the output of the EPI2ME Clone Validation workflow could be stochastic, therefore, we performed five replicates of each run. Of the 25 plasmid samples tested, 21 were consistently assembled by both assemblers, but in three of five replicate runs of Flye, only 20 plasmids assembled, demonstrating the stochastic assembly rate by Flye (Figure 2A). In contrast, Canu always assembled the same 21 plasmids, indicating consistency in assembly rate (Figure 2B). The four samples that consistently could not be assembled are all of the same plasmid at different concentrations and minipreps - it is a medium sized (12 kbp) plasmid with repeated DNA parts and is used as an AND gate in plant protoplasts (Lloyd et al., 2022). For the other plasmids that occasionally failed to be assembled by Flye, BatchA_6 failed twice (replicate 4 and 5) and BatchA_1 failed once (replicate 3). The EPI2ME Clone Validation workflow using Flye (EPI2ME-Flye) was faster per run (∼13 mins) compared to EPI2ME Clone Validation workflow using Canu (EPI2ME-Canu, ∼18 mins). In addition to assembly rate stochasticity, the assemblies produced by Flye and Canu could vary between replicate runs (Figure 2C and D). An example of this is a small deletion predicted in one replicate from EPI2ME-Flye of plasmid BatchA_3 as shown in Figure 2E. While EPI2ME-Canu did not show stochasticity at the level of assembly generation (Figure 2B), its ability to successfully reconstruct the expected plasmid sequences across five replicates did vary between runs, with various deletions in plasmid BatchA_1 predicted (Figure 2F). These results indicate that when using EPI2ME Clone Validation, multiple runs should be performed in order to be confident that the correct assembly has been generated. Overall, the successful reconstruction of the expected plasmid sequences was greater for EPI2ME-Canu than EPI2ME-Flye (Figure 2). Taken together, despite being marginally slower, EPI2ME-Canu appears to be the superior workflow given its higher rate of successful assembly of plasmids and its higher rate of assembling the expected sequence.

**Figure 2:**
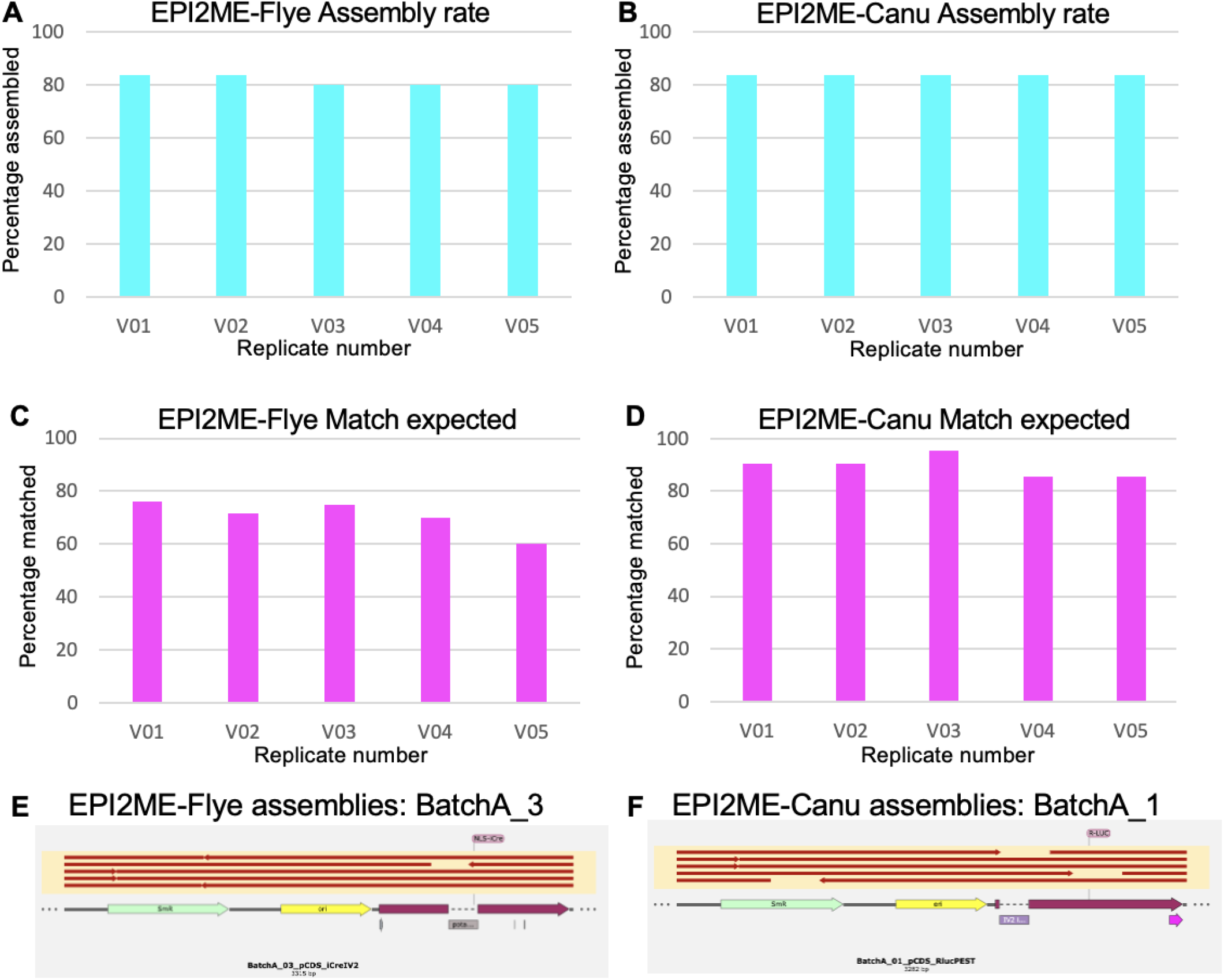
Assembly success from PICARD-seq using the EPI2ME clone validation workflow. A. The assembly rate for five replicants using EPI2ME-Flye. B. The assembly rate for five replicants using EPI2ME-Canu. C. The matching expected sequence in reference rate for five replicants using EPI2ME-Flye. D. The matching expected sequence in reference rate for five replicants using EPI2ME-Canu. E. Stochasticity in assembling BatchA_3 plasmid with EPI2ME-Flye. F. Stochasticity in assembling BatchA_1 plasmid with EPI2ME-Canu.

To better understand why some plasmids were hard to assemble for EPI2ME Clone Validation (Figure 2), we examined a dilution series of our plasmids, normalizing the concentration to molarity rather than mass (ng/µl), to account for differences in size and ensure an equivalent presence of DNA molecules per sample. Plasmids BatchB_9, _10, _11, and _12 are all the same plasmid encoding the *RUBY* coding sequence (He et al., 2020), but diluted down to different levels (2.5, 5, 10, and 20 fmol). Examining the read length distribution for these libraries indicates a very high number of reads reflecting the full-length plasmid (Figure 3A). However, the fraction of full-length reads decreases as the sample is diluted, suggesting that the less DNA per reaction the more tagmentation takes place per DNA molecule, leading to an average decrease in read length. However, even at the lowest concentration (2.5 fmol), there was a significant number of full length reads (Figure 3A). For the problematic AND gate plasmid that was not being assembled, BatchA_9, _10, _11, _12 (Lloyd et al., 2022), these samples represent a larger, more repetitive plasmid that has been diluted to a greater extent (0.5, 2.1, and 5.2 fmol). It is clear from the read distribution plots that there are very few full-length reads in each library, and no full length reads in the most diluted sample of 0.5 fmol (Figure 3B). However, despite few full length reads, there were still a significant number of long reads (≥2 kbp) in most libraries (Figure 3B), suggesting that assembly may be possible after optimising the parameters of the assembler. Therefore, we reduced the coverage requirement needed by EPI2ME Clone Validation. The default coverage parameter is 60x, so we tested it at 40x and 20x coverage for both assemblers. We found that by lowering the coverage requirement we were able to get a higher assembly rate of all plasmids, including a large AND gate plasmid (Figure 3C). Despite extensive parameter changes, a single library out of the 25 tested was recalcitrant to assembly (batchA_11). This is unsurprising given that this sample was diluted to only 0.5 fmol before sequencing (5 ng input DNA). The resulting sequencing library was depleted for full-length reads, had fewer reads total, and those reads were on average shorter than for the same plasmid at higher concentration (Figure 3B). These data suggest that a sample of plasmid that is too dilute will get over fragmented during the tagmentation step and will result in a library with too few long reads to correctly assemble the desired plasmid. Furthermore, these results indicate that the default settings for EPI2ME Clone Validation workflow are too stringent, and that by lowering the coverage parameter to 20x rather than the default 60x, more complex samples can be successfully assembled.

**Figure 3:**
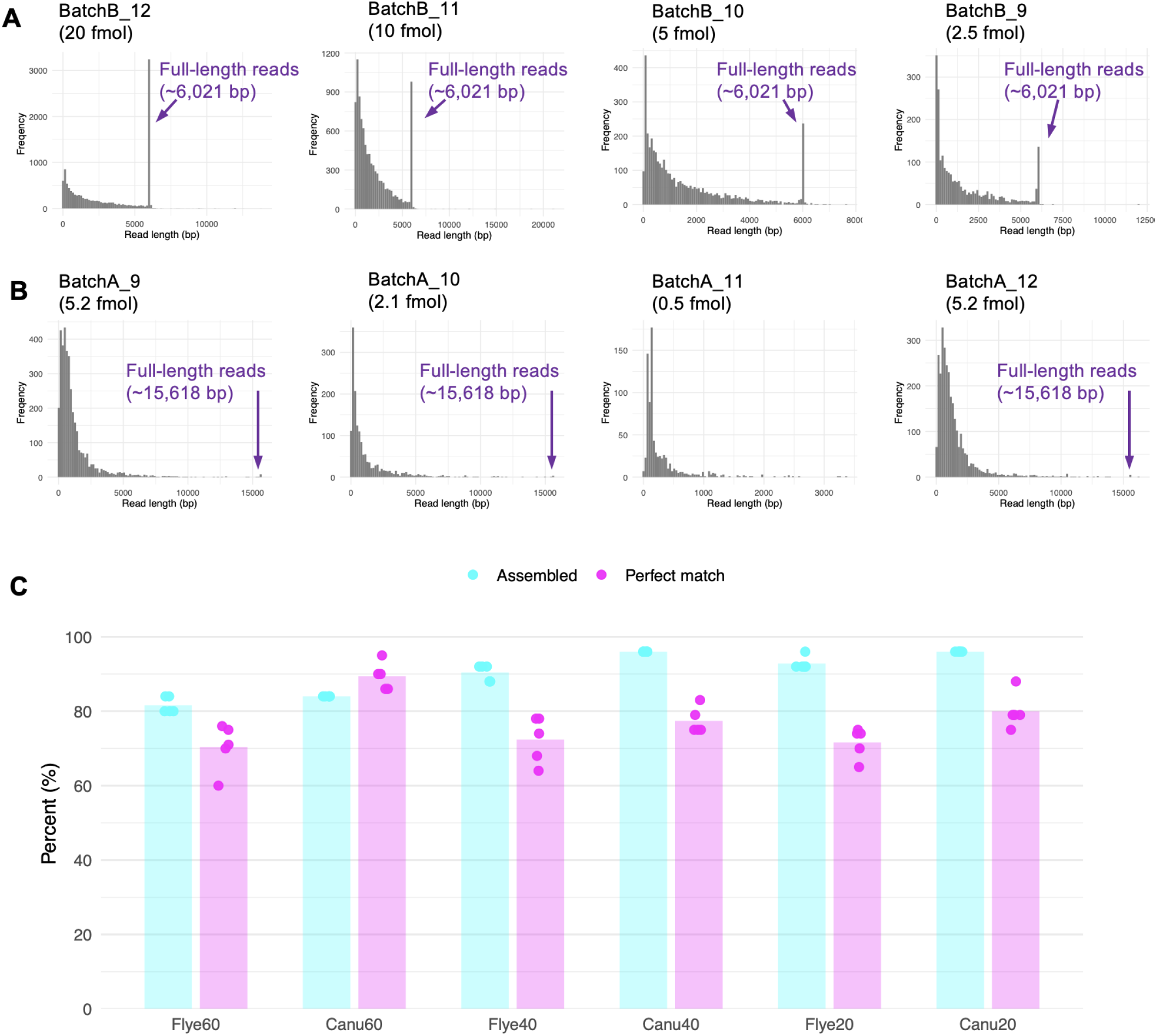
Optimization of EPI2ME clone validation workflow for PICARD-seq. A. Distribution of read lengths from samples BatchB_9-12 (pCDS_RUBY). Each sample represents the same plasmid but at a different dilution. B. Distribution of read lengths from samples BatchA_9-12 (2-input AND gate). Each sample represents the same plasmid but at a different dilution or independent plasmid extraction. C. The assembly rate and perfect match rate to reference sequences by assembler and coverage parameter in EPI2ME clone validation workflow. Percent assembled represents the number of plasmids that had an assembly out of the possible 25. The percent perfect match represents the number of plasmids that were assembled that match the expected sequence.

### Autocycler is a slower but effective assembler

To test an alternative computational pipeline to the two default assemblers provided through EPI2ME, we used the bacterial genome assembler autocycler (Wick et al., 2025). Autocycler is designed as an ensemble assembler that integrates multiple different assembly tools for genomes or plasmids, building a consensus from these multiple different construction attempts (Wick et al., 2025). Briefly, autocycler subsets the reads and runs the same tool in parallel multiple times, and then runs multiple different assembly tools in the same way to then find a consensus (Wick et al., 2025). Consequently, we noticed that this computational pipeline is significantly slower than the EPI2ME Clone Validation workflow (∼111 minutes). Despite this, the successful assembly rate was substantially higher relative to most of the EPI2ME Clone Validation runs, and the successful match rate to expected was the highest of all tested conditions (Figure 4A). These data indicate that autocycler compromises run time for a superior workflow for correct plasmid assembly. However, unlike EPI2ME Clone Validation, which integrates plasmids annotation via pLannotate (McGuffie and Barrick, 2021), this pipeline does not automatically annotate the plasmids. Nonetheless, software such as pLannotate is available independently and can be used downstream of the assembly. Overall, only one of the 24 assembled plasmids did not match the expected sequence (Figure 4A). Upon further inspection, this plasmid encodes a promoter sequence from *Arabidopsis thaliana* and contains a single base deletion relative to the expected sequence, in a large homopolymer array of A residues. Given that ONT is known to have issues determining the correct number of bases in a homopolymer (Ye et al., 2025), this is likely a mistake from the sequencing and assembly. This serves as a reminder to align and check unexpected sequence deviations to determine if they are a sequencing artefact or a real issue with the plasmid. Given that autocycler is an ensemble assembler and not all of the tools may be appropriate for plasmids, assemblers that were not appearing to benefit the pipeline were removed from the workflow in order to decrease its run time. Through creating this autocycler minimum setting, the pipeline run time decreased to only ∼81 minutes, down from ∼111 minutes. Surprisingly, one of the five runs testing this new setting was also able to correctly assemble the homopolymer region in the previously problematic plasmid (Figure 4A). Taken together, these data indicate that despite the slower speed of autocycler, it is a superior assembly pipeline compared to the EPI2ME Clone Validation workflow and is expected to provide substantial utility to many researchers.

**Figure 4:**
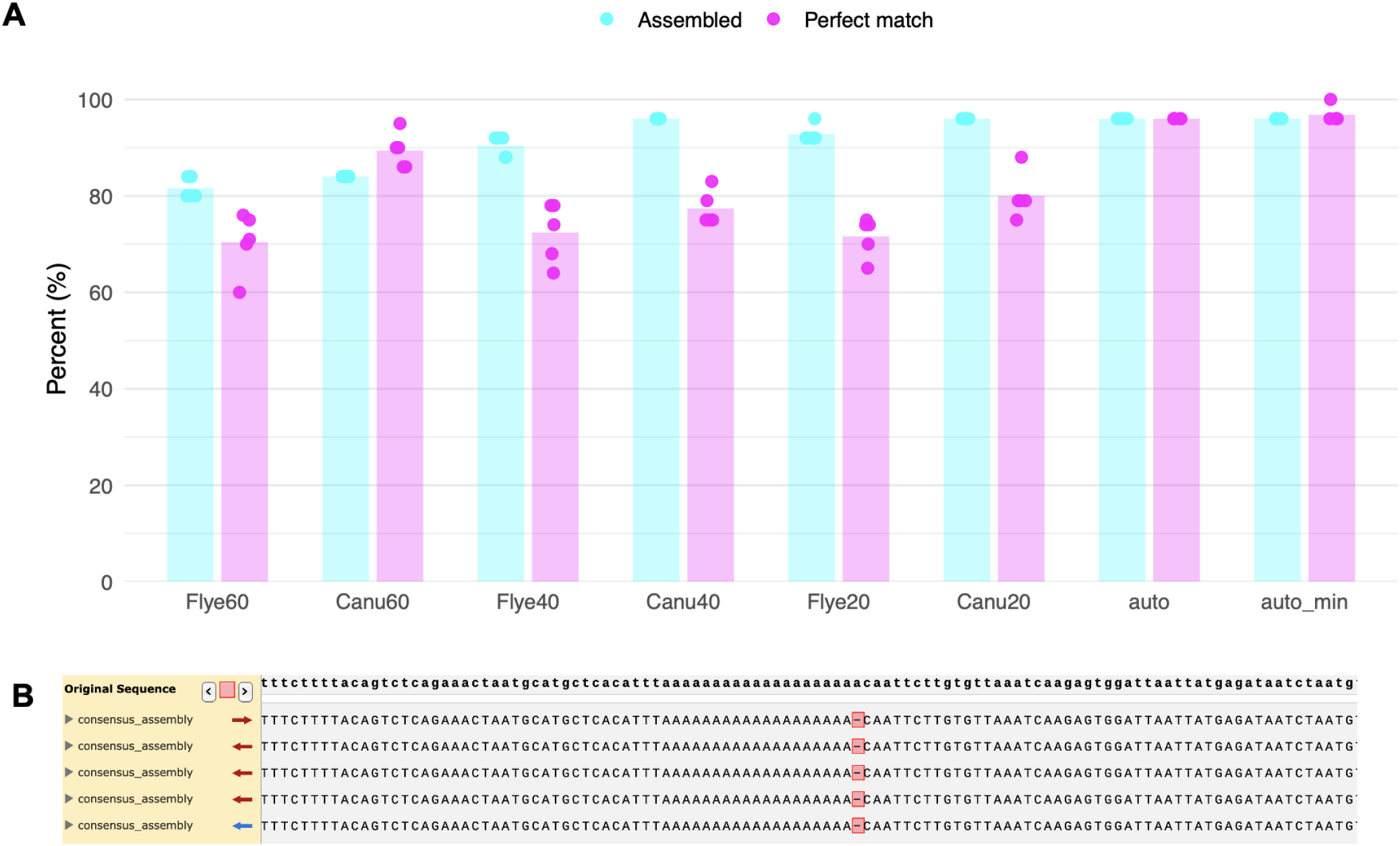
Benchmarking autocycler workflow against EPI2ME workflow. A. The assembly rate and perfect match rate to reference sequences by assembler and coverage parameter in EPI2ME clone validation workflow and autocycler workflow. Percent assembled represents the number of plasmids that had an assembly out of the possible 25. The percent perfect match represents the number of plasmids that were assembled that match the expected sequence. Autocycler = auto; Autocycler minimum = auto_min. B. Putative point mutation in plasmid BatchA_4 detected by autocycler in a homopolymer region.

### Mapping with minimap2 as an alternative validation

In addition to *de novo* assembly of plasmid sequences *in silico*, mapping the reads against a known reference sequence can also allow for determination of the correct assembly of a plasmid. Here we used minimap2 (Li, 2018; Li, 2021) to align the reads to our reference fasta file. Additionally, we developed a script that works in a conda environment to assess the quality of the mapping and generate a consensus sequence of the mapping relative to the expected reference sequence. Figure 5A highlights these values for the 25 plasmids examined. Figure 5B-E demonstrated how the mapping can vary between samples. In the case of plasmid BatchA_1, no signs of the deletions predicted by some of the EPI2ME-Canu assemblies (Figure 2F) were observed in the mapping data (Figure 5B). In the case of plasmid BatchA_11, which could not be assembled in any condition (Figure 2-4), the mapping coverage was proven to be poor and read length short, likely due to over fragmentation during library prep, given the low amount of input DNA (0.5 fmol) (Figure 5E). Furthermore, some plasmids that could be *de novo* assembled and matched the expected sequence using EPI2ME, show an imperfect match with this alternative mapping approach (Figure 5A). Therefore, this pipeline is to be used cautiously and we suggest users manually inspect the alignments in a genome browser visualisation tool rather than relying solely on the output metrics. For example, in Figure 5A and C, plasmid BatchA_7 had predicted mismatches, however, this seems to be a result of how the consensus sequence was being defined rather than an actual error in sequencing. Taken together, mapping offers an alternative and complementary way to diagnose issues with your plasmid sequencing results, in addition to the *de novo* assembly approach.

**Figure 5:**
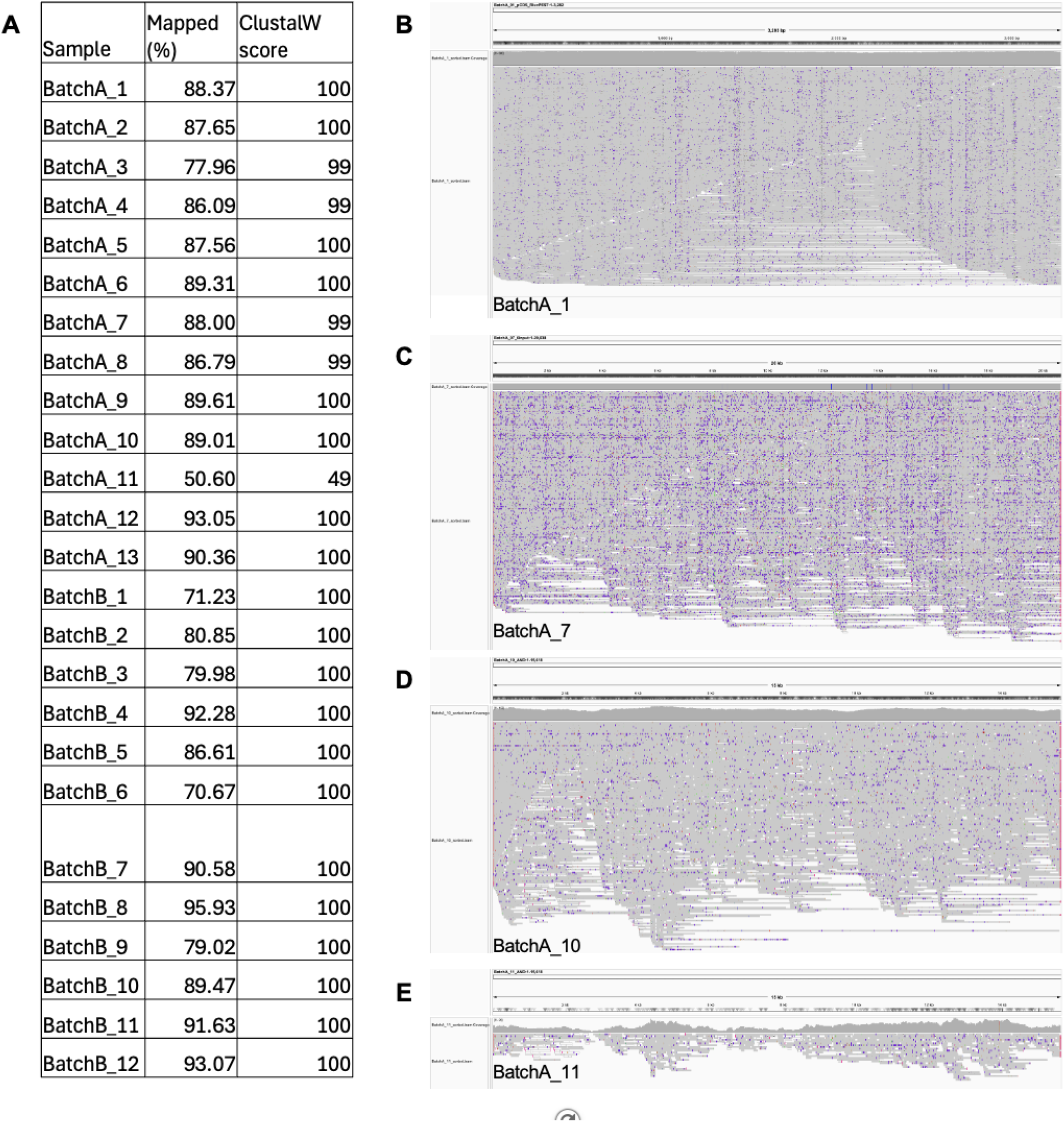
Mapping reads to expected sequence with minimap2. A. Table showing the mapping rate and the ClustalW alignment score between the consensus of alignment and the expected sequence for each plasmid. B. IGV screenshot of mapped reads to plasmid BatchA_1. C. IGV screenshot of mapped reads to plasmid BatchA_7. D. IGV screenshot of mapped reads to plasmid BatchA_10. E. IGV screenshot of mapped reads to plasmid BatchA_11.

### Rogue’s gallery of strange plasmids

Assembly of plasmids in the laboratory can yield many unexpected arrangements in the final product, and thus we strongly promote the notion that at every step of molecular cloning, sequencing is needed to validate the entire plasmid, even if it has not gone through a PCR or DNA synthesis step. Here we present a rogue’s gallery of problems that PICARD-seq has detected. For the first example, we identified an assembly containing multiple antibiotic resistant backbones. The expected sequence was a Level 2 MoClo assembly labelled the Y plasmid (Figure 6A), however, what was sequenced contained multiple antibiotic resistance genes from Level 1 (ampicillin) and Level 2 (kanamycin) MoClo constructs (Figure 6B), leading to a much larger than expected plasmid size. This sort of concatenation was unexpected but can be detected easily through sequencing with PICARD-seq.

**Figure 6:**
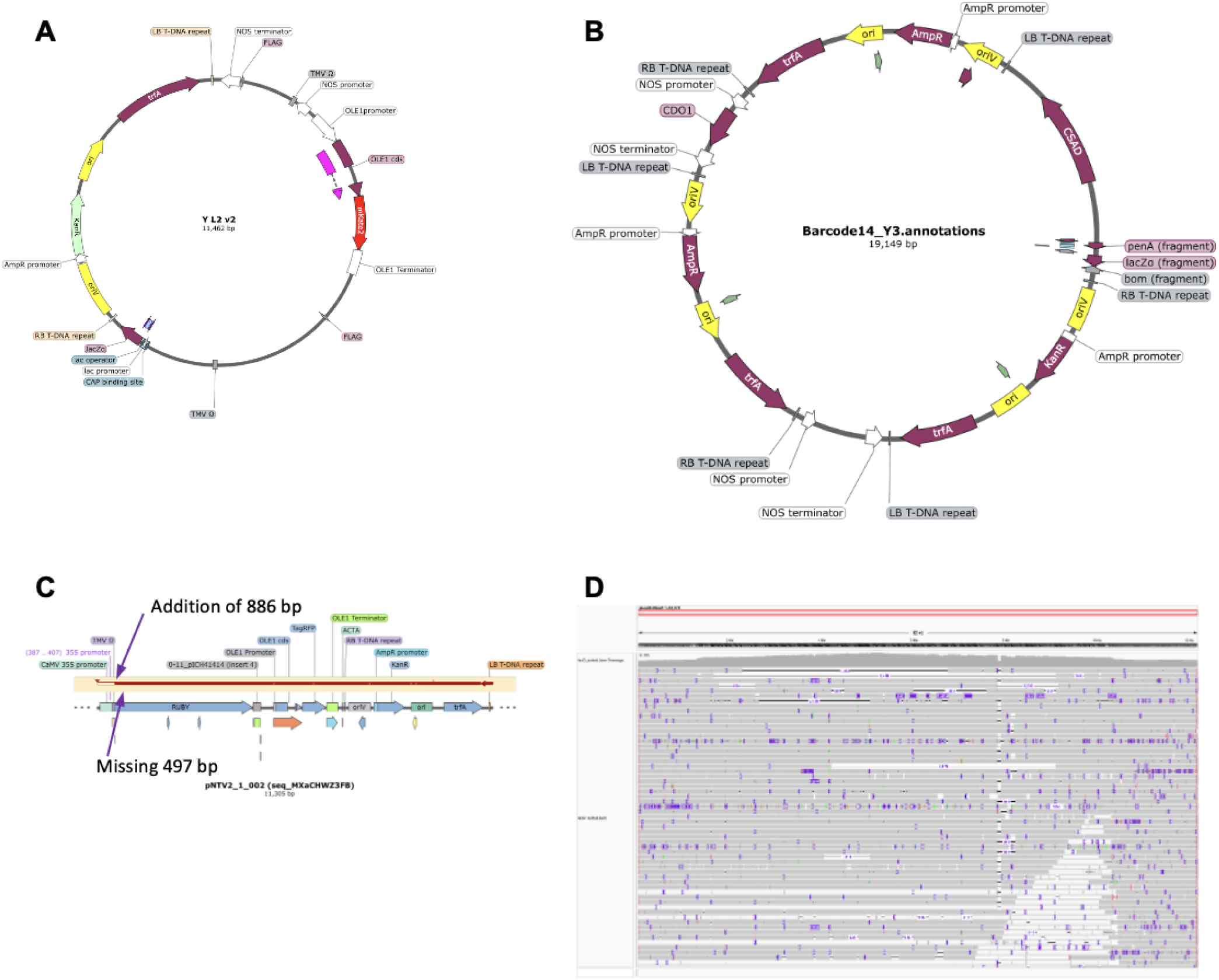
Rogues’ gallery of incorrect plasmids detected by PICARD-seq. A. Expected sequence of the Y plasmid, which encodes for a cysteine dioxygenase gene under the *NOS* promoter, a seed coat red fluorescent protein under the *OLE1* promoter, and cysteine sulfinic acid decarboxylase under the *ACT2* promoter inserted into the universal level 2 acceptor vector pAGM4673. B. Assembled plasmid for the Y plasmid from the ONT sequence run showing multiple background inserts. C. A plasmid that encodes for *RUBY* in plants. The expected promoter sequence was the *35S* (497 bp), but instead a longer sequence was identified that matched the *RbcS2B* promoter (886 bp). D. IGV genome browser view of a *lacO*-rich plasmid (pLacO-ISce1Addgene #58505). Various reads show insertions and deletions indicating a highly mixed population of plasmids.

Our next example is a construct designed to overexpress the visual reporter *RUBY* under the strong *35S* promoter in plants (Figure 6C). To our surprise, the *35S* promoter was not detected in the sequencing results (Figure 6C), however, a larger sequence was inserted in that exact same position (Figure 6C). After a BLAST search of the inserted sequence we determined it was instead the *RbcS2B* promoter. The DNA parts used for this plasmid assembly were from the Addgene kit #1000000047 (Engler et al., 2014) and it is clear that these DNA parts were mixed up when the ligation was set up. The MoClo-based assembly is particularly prone to such errors, as the standardized overhangs shared by all parts within a given position mean that an incorrectly selected part will still ligate successfully, allowing the mix-up to go undetected. Therefore, plasmid sequencing after all ligation steps is able to detect DNA part mixups that can be missed by restriction enzyme digest analysis alone. If left undetected, this mix-up would lead to unexpected results and wrongful interpretation.

For the final example, we examined a highly repetitive plasmid that contains many copies of the *lacO* sequence that is commonly used to recruit transcription factor binding (Roukos et al., 2013). After mapping the long reads to the expected sequence, the consensus matched the expected sequence (Figure 6D). However, upon closer inspection of individual reads, we identified a mixed population of reads that had large deletions and small insertions throughout the plasmid (Figure 6D). Given that ONT reads reflect single molecules (no PCR steps in library preparation), these assorted reads indicate a mixed population of rearrangements existing in a single preparation of plasmid DNA and indicate the need for sub-cloning to rectify the issue. It is likely that individual recombination events altered the size and sequence integrity of the plasmid as it has previously been reported that large repetitive plasmids are more susceptible to rearrangements (Hossain et al., 2020). Here we present strong evidence that long-read sequencing is an ideal way to detect these rearrangements and allow for better characterisation of the expected plasmid sequence. Taken together, these examples demonstrate the importance of sequencing plasmids at every step of cloning, and highlight the diversity of issues that can arise and be detected through PICARD-seq.

## Discussion

Recombinant DNA techniques that produce plasmids are a core part of biotechnology and synthetic biology. Given that many plasmids are found to contain errors (Bai et al., 2025), it is important to sequence plasmids frequently including the whole plasmid rather than just an inserted fragment region. Here we outline the use of PICARD-seq (Lloyd et al., 2026), which uses the ONT long read sequencing platform and reagents to quickly sequence entire plasmids and allow for their validation via *de novo* assembly. The advantage of this approach is that the long reads from ONT platforms are able to sequence even large repetitive plasmids. For example, we here include a 21 kbp plasmid with multiple repeated promoters and terminators that are longer than the average Illumina read length and therefore would be challenging to assemble with short reads alone. Sanger sequencing would also prove difficult and expensive to sequence the whole plasmid. Therefore, the use of ONT long read sequencing allows for an affordable and rapid diagnosis of plasmid integrity, with the typical time taken to fully sequence a set of plasmids spanning an afternoon. In this study, we have examined and compared the bioinformatics tools for analysis of the raw sequencing data. We found the ONT supported EPI2ME Clone Validation workflow to be quick, but it was highly stochastic: it varied in how many plasmids were assembled and how they were assembled between independent runs (Figure 2). We also observed that many plasmids could not be assembled with the default settings despite having many long reads in the library. We then found that by lowering the minimum coverage requirement within EPI2ME Clone Validation down to 40x or 20x, this allowed for these plasmids to be assembled correctly (Figure 3). Therefore, we suggest that if this workflow is used, the optimal parameters should be changed from the default settings (Figure 3). We then validated autocycler (Wick et al., 2025) as an alternative assembly tool. We found the assembly rate to be very high and consistent between runs, and most of the assembled plasmids matched the expected sequence (Figure 4). One limitation we found was that a single plasmid was continuously misassembled suggesting a point mutation (Figure 4). Upon closer inspection, it was identified to be an incorrectly predicted deletion of an ‘A’ residue in a homopolymer tract (Figure 4B), which is a known detection issue with nanopore sequencing (Ye et al., 2025). We then examined the utility of mapping using the alignment tool, minimap2 (Figure 5). We find that visual inspection of the mapped reads is a valuable means to identify certain DNA assembly errors, such as why some plasmids are hard to assemble and to confirm whether predicted deletions by EPI2ME Clone Validation workflow were spurious (Figure 5). Finally we have compiled a rogue’s gallery of plasmids that did not assemble as expected that we were able to identify and troubleshoot via PICARD-seq (Figure 6). This included identifying plasmids where the backbone had undergone major rearrangements relative to expected, as well as detecting the insertion of the wrong promoter sequence into a plasmid via MoClo Golden Gate cloning (Figure 6). Spotting such unexpected cloning issues would be a challenge without whole plasmid sequencing with long reads, resulting in substantial time-loss of downstream applications of the plasmid.

We see this technology as a democratising tool to allow many laboratories to be able to perform in-house plasmid sequencing at an affordable cost. Unlike other sequencing platforms, it is affordable to set up with an initial buying cost starting at $4,900 AUD ($3,150 USD) for a MinION Mk1D, plus the cost of a computer able to run it. Cost per reaction with a new flow cell is estimated to be between $4.86 and $11.18 AUD ($3.46 and $7.96 USD), depending on how many plasmids are in a run. Using old flow cells from genome sequencing projects is an effective way to further reduce this cost as the flow cell may otherwise not get fully used. The whole protocol can be performed from the isolated plasmid DNA to sequencing outcome within half a day, allowing for very rapid turnaround of results and the potential for multiple runs per week. Commercial services that offer similar whole plasmid sequencing do exist, however, in some isolated regions such as Perth, Australia, the turnaround time for results adds a significant delay. Therefore, in locations where speed of results is not possible, PICARD-seq allows laboratories to take that control into their own hands. The use of ONT long-read sequencing has already been put to good use in an automated and cost-effective way to measure recombination rates in synthetic gene circuits, demonstrating its versatility as a tool to aid synthetic biology (Greco et al., 2022). PICARD-seq allows for confidence in experimental setup by confirming plasmid identity and integrity.

The high rate of plasmid sequencing errors previously reported (Bai et al., 2025), and the examples of misassembly that we have detected (Figure 6) highlights the importance of sequencing plasmids routinely in the lab. We suggest that sequencing should take place not only after a PCR or synthesis step, but after every assembly step to identify unexpected changes to both the insert and the backbone, as well as upon receipt of any new plasmid. This will ensure that researchers are testing the expected plasmid, rather than the misassemblies that appear to be relatively common. With PICARD-seq, this opens up the possibility to rapidly and affordably validate plasmids in-house.

## Material and methods

### Mini preparation of plasmid DNA

Plasmid DNA was extracted from 1.5 ml of *E. coli* culture via the FavorPrep™ Plasmid Extraction Mini Kit (FAPDE 001-1).

### Protocol for preparing plasmids for sequencing

PICARD-seq as outlined in detail in the protocol deposited at protocols.io (Lloyd et al., 2026) was used for all plasmids. Briefly, plasmid DNA was mixed with the Tn5 enzyme with a unique barcode, allowing for the circular DNA to be linearized and uniquely barcoded for multiplexing using the Rapid Barcoding Kit 96 V14 (SQK-RBK114.96). Each reaction was then combined and bead purified before being activated by a motor protein and loaded onto the flow cell (FLO-MIN114). The flow cell was primed in accordance with standard procedure (Lloyd et al., 2026). Sequencing was then done with the software minKNOW. The flow cell was then washed for future use with EXP-WSH004 kit.

### Collection of data from sequencing run

An Apple MacBook M4 pro processor, 48 GB of RAM and 20 GPU cores, running macOS Tahoe 26.5.2 was used to analyse all of the data. This was used to run minKNOW to process the data from the MinION in real time, using super high accuracy settings (model version 5.2.0). Data was stored in POD5 and FASTQ formats.

### Plasmids for benchmarking the analysis of bioinformatics pipelines

A total of 25 plasmids with known sequences were used in this study to act as benchmarks for the assembly software (Table 1). These 25 samples have been used and sequenced using multiple platforms to ensure accuracy of the expected sequence. All sequences generated by the assembly software were compared to the expected sequence and ranked for accuracy, if they were a perfect match. Some of these plasmids represent repeats of the same construct but diluted or separately prepped to look for consistency in assembly. Some sequencing libraries were downsampled to 1000 reads to make them more representative of a standard sequencing run, rather than to sequence to excess.

### EPI2ME Clone Validation workflow

Nextflow version 24.10.5 (Ewels et al., 2020) was used for all analysis with the EPI2ME Clone Validation workflows. The wf-clone-validation v1.7.3-g229efe3 version was used for each analysis. Both Flye and Canu were used as assemblers: this choice was mutually exclusive. Additionally, the coverage level required for assembly was reduced to 40x and 20x from a default of 60x via the `assm_coverage` parameter.

### Autocycler workflow

The autocycler software (Wick et al., 2025) version 0.7.0 was installed on the above described MacBook in a custom conda environment based on advice from the developer. Modifications to the conda environment were made to allow more software to be installed on this ARM device by using an Intel version of the software. Despite this change, the assembler wtdbg could not be installed as it had not been compiled for macOS. A modified yml script was used as detailed below:

~~~
name: autocycler
channels:
   - conda-forge
   - bioconda
dependencies:
   - autocycler>=0.7.0      # https://github.com/rrwick/Autocycler
   - canu>=2.3 # https://github.com/marbl/canu
   - flye>=2.9.6            # https://github.com/mikolmogorov/Flye
# - lja>=0.2                # https://github.com/AntonBankevich/LJA
      -metamdbg>=1.4
                            # https://github.com/GaetanBenoitDev/metaMDBG
   - miniasm>=0.3           # https://github.com/lh3/miniasm
   - minimap2>=2.31         # https://github.com/lh3/minimap2
   - minipolish>=0.2.1      # https://github.com/rrwick/Minipolish
   - myloasm>=0.6.0         # https://github.com/bluenote-1577/myloasm
   - necat>=0.0.1_update20200803 # https://github.com/xiaochuanle/NECAT
   - nextdenovo>=2.5.2      # https://github.com/Nextomics/NextDenovo
   - nextpolish>=1.4.1      # https://github.com/Nextomics/NextPolish
   - plassembler>=1.8.3     # https://github.com/gbouras13/plassembler
- racon>=1.5.0              # https://github.com/lbcb-sci/racon
   - raven-assembler>=1.8.3 # https://github.com/lbcb-sci/raven
~~~

### Mapping reads to reference with minimap2

Reads for each barcode were mapped to the corresponding expected plasmid sequence using minimap2 (version 2.28-r1209) (Li, 2018; Li, 2021) with the `map-ont` preset. Alignments were sorted and indexed with SAMtools (version 1.21) (Danecek et al., 2021), and mapping statistics were recorded with `samtools flagstat`. Two consensus sequences were generated independently for each sample: one using samtools consensus, and one using ivar (version 1.4.4) consensus applied to a `samtools mpileup (-aa -A -d 5000 -Q 20)` with a frequency threshold of 0.7 and a minimum depth of 10. The ivar consensus was aligned pairwise against its reference using ClustalW (version 2.1) to obtain an alignment score and to allow inspection of any discrepancies. Intermediate SAM files and alignment guide trees were removed on completion. The complete per-sample procedure was implemented as a shell script (`minimap2_fastq_mapping.sh`), which takes a read directory, a reference FASTA and an output prefix, and writes a per-sample summary reporting total reads, mapping rate, reference and consensus lengths, and alignment score.

To apply this procedure across all plasmids, a batch wrapper (`batch_minimap2_mapping.sh`) was used. The wrapper reads a three-column sample table specifying, for each plasmid, its read directory, sample name and reference sequence, and invokes the per-sample script for each row in turn, writing the output to a separate subdirectory per sample. Samples with missing read directories or references are recorded and skipped rather than terminating the batch, and per-sample summaries are collected into a single tab-separated table of mapping and alignment statistics for downstream comparison. Both scripts are available at https://doi.org/10.5281/zenodo.22807713. Data was visualized in the Integrative Genomics Viewer (IGV) (Robinson, 2011; Thorvaldsdóttir et al., 2013).

## Data availability

Raw data, assemblies and scripts are available at https://doi.org/10.5281/zenodo.22807713.

## Acknowledgments

This work was funded by the Clifford Bradley Robertson and Gwendoline Florence Robertson fund at The University of Western Australia (UWA), UWA the School of Molecular Sciences, and the Grains Research & Development Corporation (GRDC) Mid-Career Fellowship (UWA2505-006RTX) to JPBL; the UK’s Advanced Research + Invention Agency (ARIA) Synthetic Plants programme and the Australian Research Council (ARC) Discovery Project (DP240103385) to JPBL and RL; a UWA Research Collaborative Award (2023/GR001354) and Robert and Maude Gledden Visiting Fellowship to SR and JPBL; ARC Centre of Excellence in Plants for Space (CE230100015) and National Health and Medical Research Council Investigator Grant (2035042) to RL; Australian Centre for RNA Therapeutics in Cancer and RNA Innovation Foundry support to AF; and the Centre for Crop and Disease Management, a co-investment between Curtin University and the GRDC under grant CUR00023, to JWB. HJ, PG, FL, and JYZ were supported by the UWA Research Training Program (RTP) Scholarship. HJ was supported by the Twist Bioscholar Program and ARC CoE Plants for Space HDR Scholarship.

